# Leveraging protein-protein interactions in phenotype prediction through graph neural networks

**DOI:** 10.1101/2024.08.13.605573

**Authors:** Riccardo Smeriglio, Joana Rosell-Mirmi, Petia Radeva, Jordi Abante

## Abstract

Current genotype-to-phenotype models, such as polygenic risk scores, only account for linear relationships between genotype and phenotype and ignore epistatic interactions, limiting the complexity of the diseases that can be properly characterized. Protein-protein interaction networks have the potential to improve the performance of the models. Moreover, interactions at the protein level can have profound implications in understanding the genetic etiology of diseases and, in turn, for drug development. In this article, we propose a novel approach for phenotype prediction based on graph neural networks (GNNs) that naturally incorporates existing protein interaction networks into the model. As a result, our approach can naturally discover relevant epistatic interactions. We assess the potential of this approach using simulations and comparing it to linear and other non-linear approaches. We also study the performance of the proposed GNN-based methods in predicting Alzheimer’s disease, one of the most complex neurodegenerative diseases, where our GNN approach outperform state of the art methods. In addition, we show that our proposal is able to discover critical interactions in the Alzheimer’s disease. Our findings highlight the potential of GNNs in predicting phenotypes and discovering the underlying mechanisms of complex diseases.

## I. INTRODUCTION

Single-nucleotide polymorphisms (SNPs) are genomic positions where the DNA sequence varies across populations. Through the 1000 Genomes Project, over 84.7 million SNPs have been identified [1]. Understanding the relationship between the genetic variants in these loci and phenotypes is critical for advancing our understanding on the molecular basis that underpins human disease.

The ease of obtaining vast genetic data through modern sequencing technologies has led to the widespread use of genome-wide association studies (GWAS). These studies have been incredibly valuable in detecting genetic variations linked to complex diseases such as Alzheimer’s disease (AD) [2], diabetes [3], and cancer [4]. However, GWAS typically identify linear associations, often overlooking SNP interactions, which are crucial for understanding diseases and drug development [5].

Over the past decade, the availability of those large genetic cohorts and the advancement of machine learning (ML) techniques have encouraged the interest in phenotype prediction using ML. This journey has seen the evolution from generalized linear models like Logistic Regression (LR), to more sophisticated nonlinear tree-based models, such as Random Forest (RF) or eXtreme Gradient Boosting (XGB), which have shown improvements in predictive performance [7]. In recent years, neural networks have emerged as a prominent tool for elucidating the intricate non-linear associations between genetic compositions and observable phenotypic traits. Specifically, Graph Neural Networks (GNNs) stand out in complex biological applications and have been pivotal in diverse fields like computer vision [8] and healthcare [9]. In the context of phenotype prediction, GNNs have only been recently applied showing improvements over the state-of-the-art (SOTA), showing that protein–protein interactions lead to superior results [10]. However, this work was limited to different subsets of only 101 genes previously associated with AD, divided into subsets containing at most 52 genes, and it focused on missense variants summarizing the genotype of each gene in a single dimension.

Here, we extend the set of SNPs used, encoding the genotype of each gene as a vector, and we benchmark different GNN architectures. In addition, we incorporate demographic information (age, sex) into the model, key variables in neurodegenerative diseases. Furthermore, we explore the potential of GNN approaches to unravel gene subnetworks that play a critical role in the prediction. Through simulations, we benchmark the GNN approaches against established machine learning models, including LR, multilayer perceptrons (MLPs), and tree-based methods such as RF, XGB and light Gradient Boosting Machine (LGBM). Laslty, we benchmark these approaches with data from the Alzheimer’s Disease Neuroimaging Initiative (ADNI) [11] genetic cohort of 808 subjects (Fig. 1a) alongside to clinical profiles and neuroimaging metrics. Finally, we also assess the capability of GNN approaches to discover critical epistatic interactions through GNNExplainer [12]. Our analysis provides a comprehensive picture of the behavior of GNN approaches for phenotype prediction and therapeutic target discovery in the setting of AD.

**Fig. 1:**
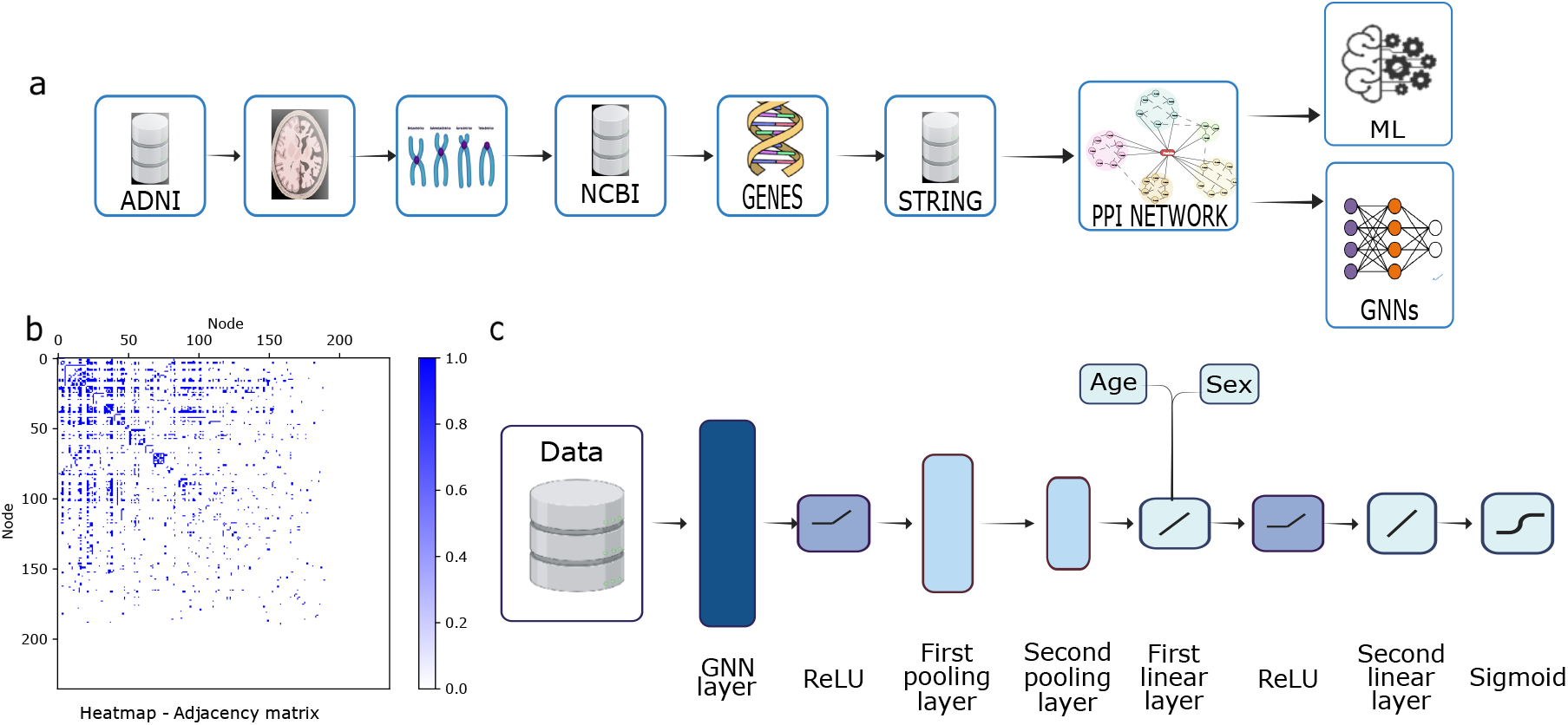
Pipeline and GNN architecture. **a**. Genomic data from AD patients are collected and encoded into a graph using a PPI network, which serves as input for the GNNs and traditional ML methods. **b**. Heatmap displaying the adjacency matrix with lower and right parts filled with zeros due to nodes not in the PPI network per STRING [6] that were not removed though they may still provide valuable insights. **c**. Both architectures comprise a GNN layer, two pooling layers, two linear layers, and a sigmoid function for class probability extraction.

## II. MATERIALS AND METHODS

### A. Graph neural network models

GNNs are neural networks that incorporate relational information between the features during inference [13]. The goal of a GNN is to learn a function of features on a graph 𝒢 = ( 𝒱,*ℰ*) [14]. A GNN takes as input, for each input graph, a feature as *x*_*i*_ for every node *i* summarized in a *N* ×*D* feature matrix *X* (*N* : number of nodes, *D*: number of input features for each node) and, a representative description of the graph structure in a matrix form, typically an adjacency matrix, *A* (see Fig. 1b).

GNN produces a node-level output *Z* (an *N* × *F* feature matrix, where *F* is the number of output features per node). Every neural network layer can then be written as a nonlinear function *H*^(*l*+1)^ = *f H*^(*l*)^, *A*, with *H*(0) = *X* and *H*(*L*) = *Z* (or *z* for graph-level outputs), *L* being the number of layers. The specific models differ only in how *f* (·, ·) is chosen and parameterized. Note that the first layer in a GNN is a multivariate function of the feature matrix, *X* and the adjacency matrix, *A* [15].

Different GNN architectures process information at various levels: graph, node, and edge. Here, we focus on Convolutional Graph Neural Networks (ConvGNNs) and Graph Attention Networks (GATs).

*1) ConvGNN Architecture:* ConvGNNs generalize grid data convolution to graph data by aggregating node features. However, they uniformly aggregate neighboring node features. A ConvGNN consists of a multi-layer graph convolutional network with the following layer-wise propagation rule:

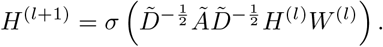

Here, *Ã* = *A* + *I*_*N*_ is the adjacency matrix of the undirected graph *G* with added self-connections. *I*_*N*_ is the identity matrix, 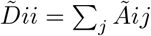, and *W* ^(*l*)^ is a layer-specific trainable weight matrix. *σ* denotes an activation function such as ReLU (ReLU(*x*) = max(0, *x*)). *H*^(*l*)^ ∈ ℝ^*N ×D*^ is the matrix of activations in the *l*th layer, where *H*^(0)^ = *X* [15].

In our setting, we use a dynamic pooling layer to select relevant nodes and a global pooling layer for overall graph patterns (Fig. 1c); two linear layers, a ReLU to introduce non-linearity and a sigmoid activation function that provides class probabilities for graph classification.

*2) GAT Architecture:* GATs introduce attentional layers, assigning different weights to nodes, thus capturing more complex graph relationships compared to ConvGNNs [16]. The GAT model used in this project is constructed with the same architecture as the ConvGNN, see in (Fig. 1c). It includes dynamic attention mechanism that allows it to focus on the most relevant and informative elements within the graph, thus, the scoring function computes a score for every edge (*j, i*), indicating the importance of the features of neighbor *j* to the node *i*, given by:

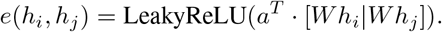

GAT computes the new representation *h*_0*i*_ for node *i* by taking a weighted average (followed by a nonlinearity, *σ*) of the transformed features of its neighbor nodes, using the normalized attention coefficients:

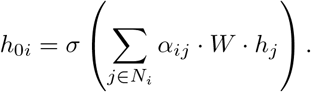

We benchmark the GNN models against LR, RF, XGB, LGBM and MLP. These models were implemented using `scikit-learn` [17], whereas the GNN approaches were implemented using `Pytorch` [18]. In addition, for each GNN model, we conducted a hyperparameter search on the number of hidden channels of the ConvGNN and heads on the GAT, considering for both 8, 16, and 32, and the number of GNN layers, considering 1, 2 or 3.

### B. Genotype Encoding

To train the model, we need to transform the genotype information into numerical values. Genotype information is usually encoded into 0 and 1’s, with the former referring to the reference allele (most common allele in the population) and the latter to an alternative allele (any other genetic variant). Following standard practice, in each position we take the mean between both chromosomes, which we assign a numeric value of 0 if the SNP represents the reference allele and 1, otherwise (Fig. 2).

**Fig. 2:**
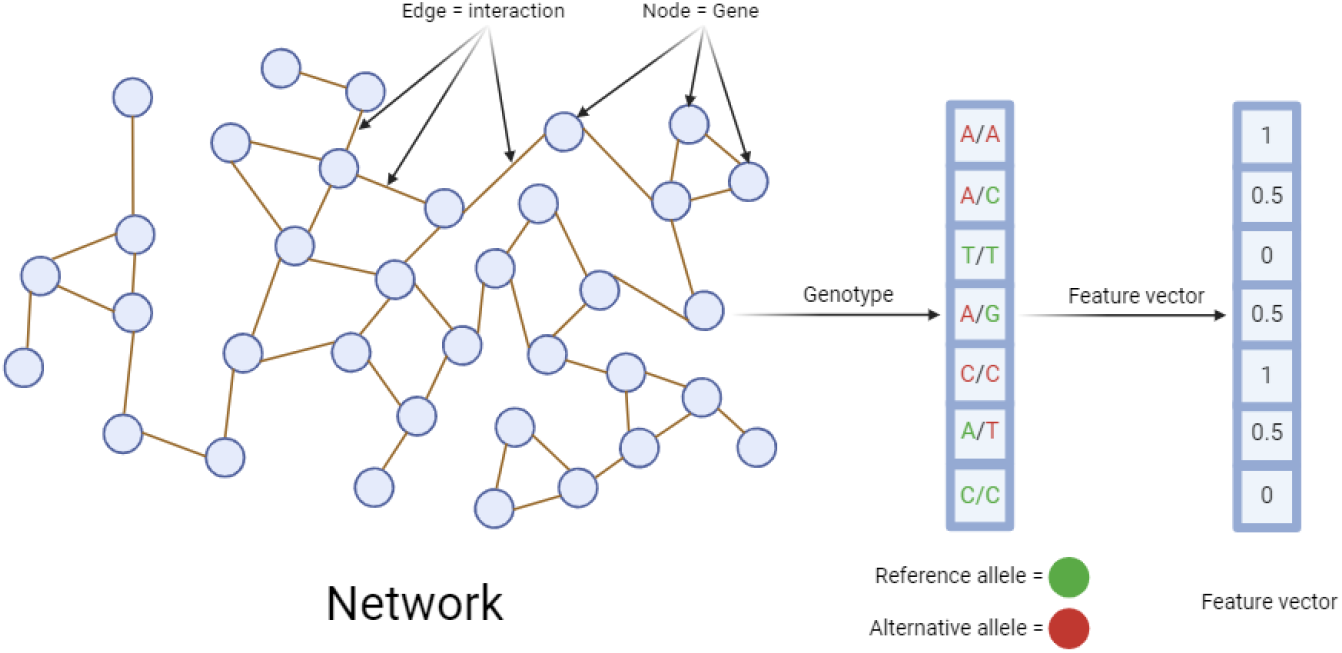
Encoding the genetic information. We assign a numeric value to each SNP in each patient by giving a 0 to those SNPs with the reference allele present in both chromosomes; a value of 0.5, when an alternative allele is present and a value of 1, when both alleles are alternative.

To fully define the graphs 𝒢 = (𝒱,*ℰ*) used to train our GNN models, we also require an adjacency matrix that defines *ℰ*. Since we want to leverage previous biological knowledge, we construct such matrix from PPI networks. In this case, the nodes, 𝒱, and the edges, *ℰ*, represent genes and gene interactions, respectively. To extract these interactions, we queried the STRING database [6]. In these graphs, each node is composed of a feature vector containing the encoded genotype of the SNPs found in the corresponding gene in the order, they are found in the gene. Multiple SNPs can be present in a given gene, resulting in different feature dimensions across nodes. To address this, we zero-padded all feature vectors to the maximum number of SNPs in a single gene. This results in a set of feature vectors with the same dimension.

### C. Simulated data

To study the effect of interactions in the performance of the models in a controlled setting, we consider two different scenarios in our simulations. First, we consider the case where the phenotype (AD diagnosis) is driven by an interaction between the genotype of APOE, a gene that plays a major role in AD [19, 20], and the age of the subject (scenario A). In this case, variants that differ from the reference allele promote the disease at a faster rate. Second, we consider the case where the phenotype is driven by interactions between APOE, Apolipoprotein B (APOB) and low-density lipoprotein receptor (LDLR) genes, known to interact with APOE [21], as well as age (scenario B). In this case, the rate at which the phenotype is developed is faster only when the disease-linked variants appear together. In both cases, we study the performance of the models for 250, 500, 750, and 1,000 simulated subjects balancing the phenotype.

#### 1. Scenario A - No interactions

We simulate the case where the fifteen SNPs contained in APOE are the ones driving the AD phenotype, with the odds ratio increasing proportionally as a function of age. Thus, the logit function is given by:

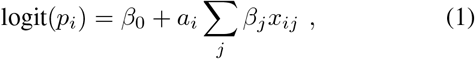

where *a*_*i*_ represents the normalized age of the *i*-th subject, and *x*_*ij*_ represents the value of the *j*-th risk-associated SNP for the *i*-th simulated subject.

#### 2. Scenario B - Interactions

In order to simulate the scenario where the interaction between multiple SNPs drive the AD phenotype, we use a LR model as in scenario A. In this case, however, SNPs related to APOE (15 SNPs), APOB (1 SNP), and LDLR (12 SNPs) were extracted from the real dataset, resulting in a total of 207 interactions when considering pairs of SNPs belonging to different genes. Thus, in this case the logit probability is given by:

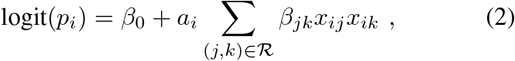

where ℛ is the set of interactions considered.

### D. ADNI data

The ADNI database is a comprehensive and long-term research study aimed at developing biomarkers for the early detection and monitoring of this neurodegenerative disease [11]. It contains genetic data for 808 patients between 50 and 91 years of age. Due to computational limitations, here we focus on chromosome 19, consisting of 484,583 SNPs, wherein the APOE gene is located.

For each patient, we retrieve values of Florbetapir F18 (AV45) or Pittsburgh Compound-B (PiB) standardized uptake value ratio (SUVR) from their last visit, the age at their last visit and their biological sex. PiB is a positron emission tomography (PET) imaging tracer that is utilized to detect and visualize the presence of amyloid deposits in the brain [22]. AV45 is a radiotracer used in amyloid PET imaging for AD research and clinical trials [23], being the amyloid PET status associated with the presence of amyloid pathology in the brain, which is one of the key features of AD [24].

To label the data, the positive or negative amyloid PET status is assigned for each subject’s last visit using a threshold of 1.27 for PiB and a threshold of 1.11 for AV45 SUVR values [10]. Thus, patients with AV45 values above the threshold are classified as class 1, indicating a positive amyloid PET status and patients below the threshold are classified as class 0. For patients without AV45 values, the same classification process is applied using the PiB value. The patients without information about PiB and AV45 SUVR are not included in the dataset, reducing the number of patients to 668.

SNPs that do not encode a specific gene according to the NCBI [25] and the GeneCards [26] databases are excluded. We have also removed SNPs with a high percentage of missing values across subjects, resulting in a dataset with 668 patients, 517 SNPs overlapping 236 genes. Regarding biological sex and age, the former is codified with a binary variable (female: 0, male: 1) and the latter normalized using a min-max scaling. To build the gene network, we use PPIs information from STRING [6], including both direct (physical) and indirect (functional) associations, and represent it using a network. The network used for our GNN models comprises 236 nodes with 15 features each and 1,143 undirected edges (Fig. 1b). Given the interactions are assigned a score indicating their confidence level, we chose to employ three distinct thresholds on the PPI scores provided by STRING (0, 0.5, and 0.75), resulting in three different networks with 1,143, 798, and 302 edges in each case.

The real GNN dataset is randomly split into a training and a validation set. The training set is composed of 260 patients from class amyloid PET positive and 260 patients from class amyloid PET negative, resulting in a total of 520 patients. The remaining patients are used for the validation set, consisting of 128 elements from class amyloid PET positive and 20 elements from class amyloid PET negative. To evaluate the robustness of the classifiers, this process is repeated 10 times, resulting in 10 different training and validation sets. In addition, to evaluate the importance of the interactions in the predictive performance of the ConvGNN approaches, we also consider a ConvGNN approach where the adjacency matrix is randomly permuted.

## III. RESULTS

Here we present the simulations and real data results (Fig. 3). To benchmark the considered methods, we use balanced accuracy (BA) computed as:

**Fig. 3:**
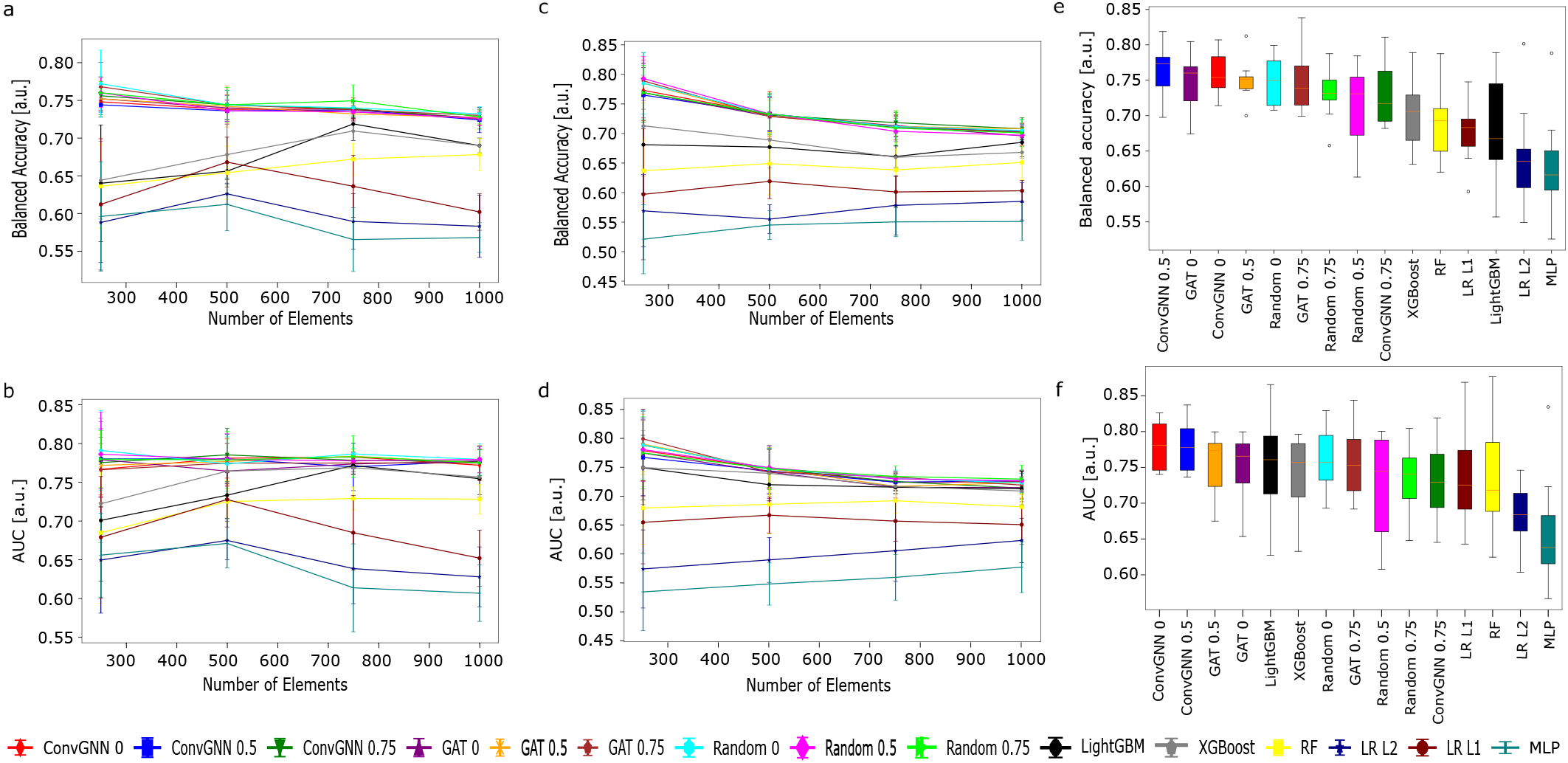
Simulations and real data results. **a**. Balanced accuracy in scenario A as the dataset size increases. **b**. AUC in scenario A as the dataset size increases. **c**. Balanced accuracy in scenario B as the dataset size increases. **d**. AUC in scenario B as the dataset size increases. **e**. Balanced accuracy in the real data analysis. **f**. AUC in the real data analysis. In (a-d) the median is shown with the bars representing an interval with ±std. In (e-f), the orange line shows the median, the box shows the interquartile range (Q3-Q1), and the whiskers extend the Q1 and Q3 by ±1.5·IQR.

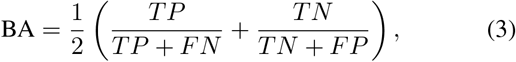

where TP represents the *true positives*, TN represents the *true negatives*, FP represents the *false positives* and FN represents the *false negatives*. In addition, we consider the area under the Receiver Operating Characteristic (ROC) curve (AUC).

### A. Simulations

Each simulated dataset is divided into five different groups of balanced elements to apply a five-fold method for training and validation of all the models for a total of 20 training and validation for each model.

#### 1) Scenario A

When no interactions are present in the simulated data, GNN approaches consistently outperform the rest of models in terms of balanced accuracy and AUC (Fig. 3a and 3b). Notably, the performance of GNN approaches does not seem to significantly improve with an increasing number of elements used for training, suggesting there is no further room for improvement. Regarding the linear approaches, the *L*_1_ regularized LR performs better than its *L*_2_ regularized counterpart, consistent with the fact that *L*_1_ regularization is more effective in terms of feature selection. Both linear models, however, fall behind the tree-based models and GNNs. Among the former, XGB stands out as the best-performing one, followed by LGBM and RF, consistent with previous observations. Interestingly, the MLP approach performs the worst in this scenario.

#### 2) Scenario B

In the presence of interactions, GNNs consistently outperform other models in terms of both AUC and BA, as illustrated in Fig. 3c and 3d. This demonstrates their ability to handle complex interactions effectively. In these scenarios, the three base methods outshine the linear methods. Notably, the performance of XGB and LGBM, in terms of AUC, is comparable to that of the GNNs. Among the linear methods, the LR with *L*_1_ regularization surpasses the LR with *L*_2_ regularization, again highlighting the implicit feature selection in *L*_1_ regularization.

### B. ADNI data

Each model was evaluated training with each of the Training set - validation set couple, for a total of 10 training and validation for each model. When benchmarking the different approaches, GNN-based models continue to show the highest BA (Fig. 3e and 3f). The ConvGNN model achieve the best performance overall when setting a threshold of 0.5 for the interaction confidence 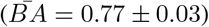, followed by the model using a GAT layer with a threshold of 0 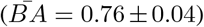 and the ConvGNN model with a threshold of 0 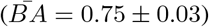. Among the ML models, the XGB model stands out as the best performer 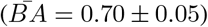, making it the only ML model surpassing 70% BA, closely followed by the RF model 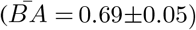 and the *LR*_*L*1_ model at 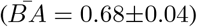. Lastly, the LGBM 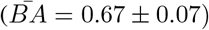, *LR*_*L*2_ 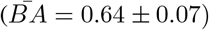, and MLP 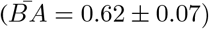 models lagged behind.

The observed differences were statistically significant when comparing, using the *Wilcoxon test*, all the GNNs with all the ML models in terms of *BA*. Exceptions include the ConvGNN with a threshold of 0.75, which did not show statistically significant differences when compared to XGB and LGBM. Additionally, the ConvGNN with the randomly permuted adjacency matrix and a threshold at 0.5 did not exhibit statistically significant differences when compared to all tree-based methods and LR with *L*_1_.

### C. GNN interpretability

One of the most interesting features about GNN approaches is the possibility to study which genes and interactions are driving the prediction of the model. To that end, we used the GNNExplainer [12] to the model that showed the highest *BA*. GNNExplainer identifies key nodes and edges in the graph, pinpointing a compact subgraph crucial for making a given prediction.

Using GNNExplainer, we extracted the most significant subgraph for each patient in the validation set (Fig. 4). This approach enabled us to identify thirteen key genes and two PPIs across all subgraphs. Interestingly, nine out of these, being CYP2S1, LDLR, XRCC1, CD209, APOE, SIRT6, DNMT1, NOTCH3, and INSR, have been previously implicated in AD [10, 21, 27, 28, 29, 30, 31, 32, 33] whereas the other four genes, RETN, PLA2G4C, AMH, and KLK3, have not. The GNNExplainer identified two significant interactions: (i) between APOE and LDLR, and (ii) between RETN and INSR. The relevance of the APOE gene to AD is already established [19, 20], but its interaction with LDLR is especially intriguing. Recent studies have shown that overexpression of LDLR in tauopathy mice significantly reduces APOE levels in the brain, ameliorating both tau pathology and neurodegeneration [21]. However, there are no studies describing the interaction between RETN and INSR in the context of AD.

**Fig. 4:**
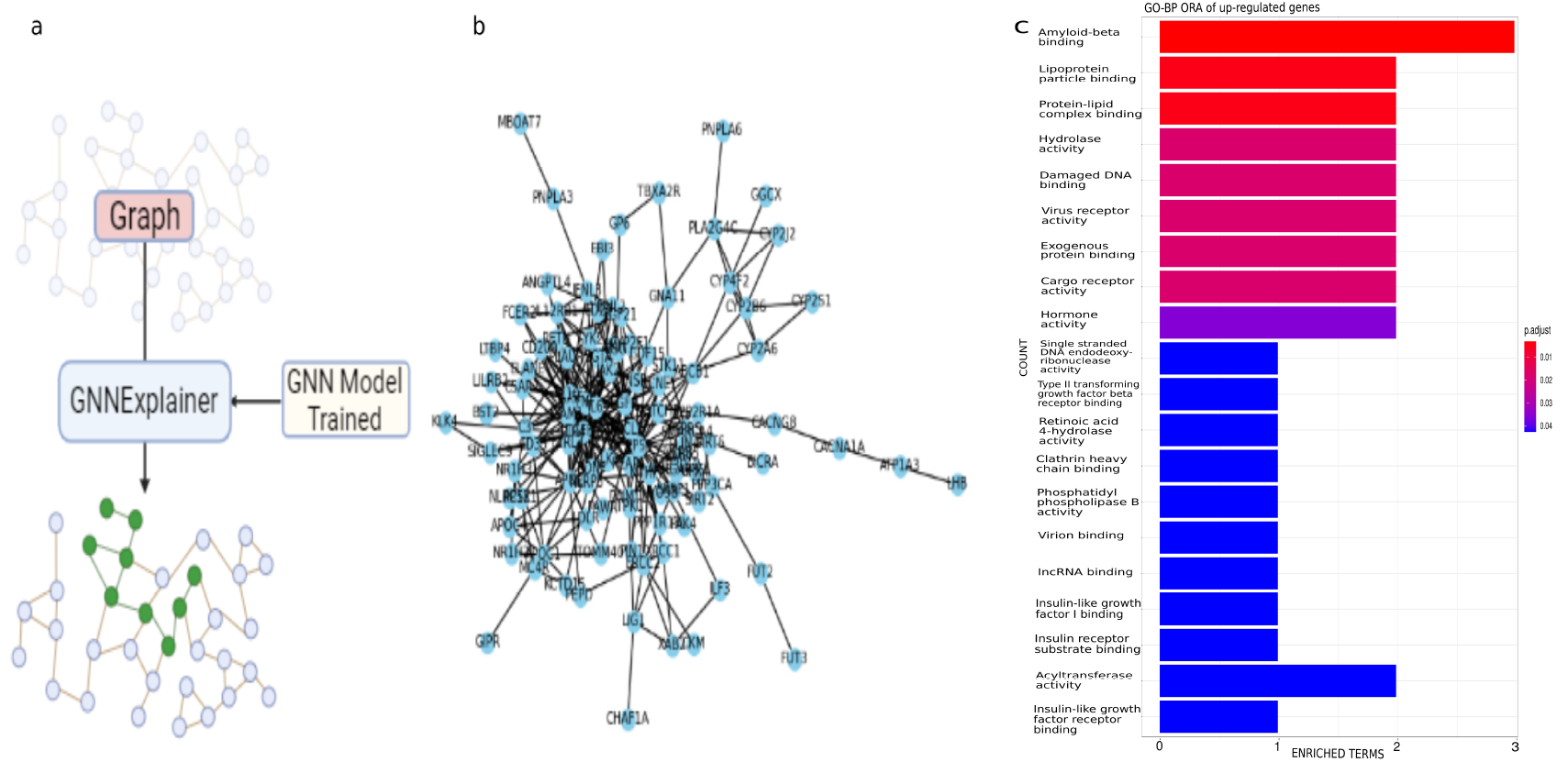
GNNExplainer explained **a**. Schematic functioning of GNNExplainer: from the input graph, and using the trained GNN model, this tool is capable of gathering the SNPs and interaction in the data that drives into the final output, for each patient identifies key nodes and edges in the graph. **b**. Example of a significant subgraph extracted using GNNExplainer. **c**. The enriched Gene Ontology terms from our analysis arranged by adjusted p-value. Interestingly, the top 3 include amyloid beta binding, lipoprotein particle binding, and protein-lipid complex hydrolase activity.

Furthermore, we did an enrichment analysis to those thirteen genes obtained with the GNNExplainer, to seek for the genes over-represented to further strengthen the explainability of our discoveries. Among the list of identified factors (Fig. 4c), the following three were the most statistically significant: amyloidbeta (A*β*) binding (p<0.001), lipoprotein particle binding (p<0.001), and protein-lipid complex binding (p<0.001).

Understanding the role of impaired movement of lipoprotein particles in the brain’s extracellular space sheds light on AD pathophysiology. Degeneration triggers A*β* release, which leads to microglial reactions and neuronal cholesterol deficiency-induced tau accumulation, thereby presenting new avenues for therapeutic exploration. Moreover, the use of radiotracers in PET imaging, like AV-45, serves as a pivotal biomarker in contemporary AD diagnosis, facilitating precise detection and quantification of A*β* deposition in vivo [21, 22, 34].

## IV. DISCUSSION

In our simulations, for small sample sizes, we observed slight differences between the GNN architectures considered which disappeared for larger sample sizes, with all the GNN methods attaining a BA*>* 0.70. While it is natural to anticipate better performance from models with larger training datasets, we observed a decrease in performance as the number of elements increased in both scenarios. This raises questions about the scalability and data efficiency of these models, which we aim to explore further in future studies. On the other hand, the performance of RF, LGBM, and XGB improved as the training dataset grew in size as expected, particularly in Scenario A. Lastly, the MLP shows the worst performance in all scenarios tested, with all models outperforming it. We attribute this to the fact that the MLP approach probably requires a larger dataset. Surprisingly, however, the GNN approaches do not seem to require as much data as one would expect, surpassing simpler models, as LR and tree-based models with only 250 subjects available.

The results obtained through our benchmark on the ADNI database are consistent with that of our simulations. GNN approaches perform the best across all considered models, highlighting the effectiveness of GNN approaches to predict AD diagnosis. This indicates that GNNs are capable of capturing the complex relationships between genetic data and the PET amyloid status of an individual. In contrast, the MLP model exhibited the worst performance (Fig. 3e and 3f), which we attribute to the small number of elements provided to the network. All GNN models displayed comparable performance (Fig. 3e and 3f).

Notably, the median performance in terms of BA and AUC remained unaffected when comparing ConvGNNs using the original versus the permuted adjacency matrix. Nevertheless, the variance in the models with permuted matrices was higher compared to the non-permuted models. We attribute this to the fact that some truly important interactions might be missing and might be added when permuting the adjacency matrix, whereas we attribute the latter to the deletion of important interactions. ConvGNN models with thresholds of 0 and 0.5 are notably superior in terms of BA when compared to their permuted matrix counterparts. Specifically, the ConvGNN model with a threshold of 0 showed a statistically significant improvement over the permuted model with thresholds of 0.5 (*p* = 0.006) and 0.75 (*p* = 0.027), and even outperformed the ConvGNN with a threshold of 0.75 (*p* = 0.037). Similarly, the ConvGNN model with a threshold of 0.5 was superior to the permuted model with thresholds of 0.5 (*p* = 0.020) and 0.75 (*p* = 0.037), and also demonstrated better performance than the ConvGNN with a threshold of 0.75 (*p* = 0.019).

Regarding the architecture of the GNN approaches, the model with the best results in both BA and AUC was the ConvGNN, with thresholds 0 and 0.5, respectively (Fig. 3e and 3f). The GAT models followed the ConvGNN model in performance probably due to the size of the model, as the GAT model contains more parameters than the ConvGNN. Furthermore, ConvGNN 0.75 was the worst performer among the graph models, as well as Random 0.5 and Random 0.75, suggesting that some edges with low confidence are important in predicting AD diagnosis. Nevertheless, here we did not find significant differences in the performance as a function of the confidence level threshold applied to the PPI network.

The approach introduced in Hernandez et al. 2022 [10] differs in many aspects to the proposed, as previously discussed, which we believe offers some advantages, resulting in an improvement of 10% in AUC (68% vs. 78%; Fig. 3). We attribute this improvement (i) to the use of a larger number of genes (at most 52 vs 236 genes), and (ii) to the fact that the features used in [10] consist of the number of missense variants in a given gene, as opposed to having each node be a numeric vector representing encoding the entire genotype, losing critical information about specific epistatic interactions, for example. Moreover, in our approach, we extended the data using demographic informations (age and sex) into the model, key variables in neurodegenerative diseases.

Finally, we have also shown how the subgraphs identified using GNNExplainer recapitulate genes that have been previously involved in AD pathogenesis. Our approach also identified an interaction between APOE and LDLR as being important in the diagnosis of AD. This interaction was recently described experimentally [21], showing the potential of our model to discover these interactions *in silico*. In addition, our model identified four genes which appear to be influencing the prediction, as well as an additional subnetwork that warrants further study. To enhance the explainability of our findings, we applied gene-ontology analysis, highlighting the significance of the identified genes in amyloid-beta binding, lipoprotein particle binding, and protein-lipid complex binding, factors previously linked to AD [21, 22, 34]. Importantly, the proposed approach provides insights into the specific subnetworks driving the diagnosis per individual, showing the potential of the approach for *in silico* precision medicine.

## V. CONCLUSIONS

Here, we studied the performance of GNN for phenotype prediction and benchmarked them against SOTA methods, outperforming them even with small training datasets. A salient application of the proposed models is their potential to empower clinicians in diagnosing a patient’s PET amyloid status, relying solely on genetic data. In addition, the proposed approach includes age and gender as features used in the prediction, allowing them to predict the likelihood of younger individuals being diagnosed with AD in the future. As a result, this predictive model could optimize primary prevention strategies even before the disease manifests. Lastly, the proposed approach provides a novel way of discovering risk-associated variants and epistatic interactions at a subject level. We showed how using this approach, we can rediscover known risk-associated variants and experimentally validated interactions, and we provided four additional genes and a novel interaction warranting further study.

## ACKNOWLEDGMENTS

Data collection and sharing for this project was funded by the Alzheimer’s Disease Neuroimaging Initiative (ADNI) (National Institutes of Health Grant U01AG024904) and DOD ADNI (Department of Defense award number W81XWH-12-2-0012). As such, the investigators within the ADNI contributed to the design and implementation of ADNI and/or provided data but did not participate in analysis or writing of this report. This study was supported by grants from the Instituto de Salud Carlos III, Ministerio de Ciencia e Innovación and European Regional Development Fund (ERDF A way of making Europe) (Red de Terapias Avanzadas, RD21/0017/0020); European Union NextGenerationEU/PRTR; Generalitat de Catalunya (2021 SGR 01094); and “la Caixa” Foundation under the grant agreement LCF/PR/HR21-00622”, Spain. This publication is also part of the project PNRR-NGEU which has received funding from the MUR–DM 118/2023. Lastly, we would also like to thank Bhalaji Nagarajan and Jesús Molina Rodríguez De Vera for their insightful suggestions and feedback.

## Notes

### Competing Interest Statement

The authors have declared no competing interest.

### Summary of Updates

typos changes and superfluous info deleted

